# *Mantoniella Beaufortii* and *Mantoniella Baffinensis* sp. nov. (Mamiellales, Mamiellophyceae), Two New Green Algal Species from the High Arctic

**DOI:** 10.1101/506915

**Authors:** Sheree Yau, Adriana Lopes dos Santos, Wenche Eikrem, Catherine Gérikas Ribero, Priscilla Gourvil, Sergio Balzano, Marie-Line Escande, Hervé Moreau, Daniel Vaulot

## Abstract

Members of the class Mamiellophyceae comprise species that can dominate picophytoplankton diversity in polar waters. Yet polar species are often morphologically indistinguishable from temperate species, although clearly separated by molecular features. Here we examine four Mamiellophyceae strains from the Canadian Arctic. The 18S rRNA and Internal Transcribed Spacer 2 (ITS2) gene phylogeny place these strains within the family Mamiellaceae (Mamiellales, Mamiellophyceae) in two separate clades of the genus *Mantoniella*. ITS2 synapomorphies support their placement as two new species, *Mantoniella beaufortii* and *Mantoniella baffinensis*. Both species have round green cells with diameter between 3–5 μm, one long flagellum and a short reduced flagellum (~1 μm) and are covered by spiderweb-like scales, making both species similar to other *Mantoniella* species. Morphologically, *M. beaufortii* and *M. baffinensis* are most similar to the cosmopolitan *M. squamata* with only minor differences in scale structure distinguishing them. Screening of global marine metabarcoding datasets indicates *M*. *beaufortii* has only been recorded in seawater and sea ice samples from the Arctic while no environmental barcode matches *M. baffinensis*. Like other Mamiellophyceae genera that have distinct polar and temperate species, the polar distribution of these new species suggests they are cold or ice-adapted *Mantoniella* species.

## Introduction

Over the last decades the taxonomy of the green algae has gone through a profound reorganization. The class Prasinophyceae, initially defined as scaly flagellates (Throndsen and Moestrup 1988), has been rearranged into many new classes such as the Chlorodendrophyceae, Chloropicophyceae and Mamiellophyceae (Massjuk 2006, Marin and Melkonian 2010, Lopes dos Santos et al. 2017b) leading to the name Prasinophyceae to be abandoned. The Mamiellophyceae are ecologically successful and particularly dominant in marine coastal waters (Lopes dos Santos et al. 2017a, Tragin and Vaulot 2018). The first scaled species of Mamiellophyceae observed were *Mantoniella squamata* (Manton et Parke) Desikachary (originally *Micromonas squamata*), in 1960 (Manton and Parke 1960), and *Mamiella gilva* (Parke et Rayns) Moestrup (originally *Nephroselmis gilva*), in 1964 (Moestrup 1984). Moestrup (1984) erected the family Mamiellaceae, which included *Mantoniella* and *Mamiella*, with *Mamiella gilva* designated as the type species. The class Mamiellophyceae comprises three orders: the Monomastigales, with one freshwater genus *Monomastix*; Dolichomastigales, with two genera *Crustomastix* and *Dolichomastix*; and the Mamiellales, which currently comprises five genera *Bathycoccus, Mamiella, Mantoniella, Micromonas* and *Ostreococcus*. As these genera are morphologically heterogeneous, with *Micromonas* and *Ostreococcus* lacking scales and *Bathycoccus* and *Ostreococcus* lacking flagella, the monophyly of Mamiellophyceae was established based on nuclear and plastid rRNA sequence and secondary structure analyses (Marin and Melkonian 2010).

Molecular analyses of the Mamiellophyceae have permitted the description of otherwise morphologically indistinguishable cryptic species. For example, wide genetic diversity has been shown to exist between morphologically identical *Ostreococcus* species where less than 1% difference in the 18S rRNA gene corresponds to up to 30% of variation in orthologous protein coding sequences (Palenik et al. 2007, Piganeau et al. 2011). From an early stage, 18S rRNA defined clades of *Micromonas* and *Ostreococcus* were observed to correspond to distinct geographic distributions, suggesting their genetic variation reflects adaptations to ecological niches (Rodríguez et al. 2005, Foulon et al. 2008), and that these clades represent distinct species. *Ostreococcus* is divided into rare species restricted to estuarine (*O. mediterraneus*) and coastal environments (O. *tauri*), as well as more abundant oceanic species (O. *lucimarinus* and clade B) (Demir-Hilton et al. 2011, Treusch et al. 2012, Hu et al. 2016, Simmons et al. 2016). In *Micromonas*, the species *M. polaris* appears to be specially adapted to polar environments, while the other species are restricted to lower latitudes (Not et al. 2005, Lovejoy et al. 2007, Balzano et al. 2012b, Simon et al. 2017). Similarly, in *Mantoniella, M. antarctica* was described from the Antarctic while *M. squamata* is cosmopolitan (Marchant et al. 1989).

Three picophytoplanktonic strains (RCC2285, RCC2288 and RCC2497) were isolated in the Canadian Arctic from mesophilic surface water sampled at two sites in the Beaufort Sea in the summer of 2009 as part of the MALINA cruise (Balzano et al. 2012a). A fourth strain was subsequently isolated from sea ice collected in Baffin Bay in the spring as part of the Green Edge project. We performed a combination of molecular, morphological and pigment characterization of these isolates, which we propose to constitute two novel *Mantoniella* species, *M. beaufortii* and *M. baffinensis*, restricted to polar environments.

## Methods

### Culture conditions

Strains RCC2285, RCC2288, and RCC2497 were isolated from seawater collected at two sites (70°30’N, 135°30’W and 70°34’N, 145°24W) in the Beaufort Sea in the summer of 2009 as part of the MALINA cruise as described previously (Balzano et al. 2012a). Strain RCC5418 was isolated from the Green Edge project Ice Camp (http://www.greenedgeproject.info/), a sampling site on the sea ice near the village of Qikiqtarjuaq (67°28.784N, 63°47.372W). Sampling was conducted between 20 April and 27 July, 2016, beginning in completely snow covered conditions followed by bare ice and ending when the ice was broken out. Sea ice from 23 May 2016 was melted overnight and 200 ml was gravity filtered (Sartorius filtration system) through 3 μm pore size polycarbonate filters (Millipore Isopore membrane, 47 mm). 500 μL of filtrate was enriched by addition to 15 ml of L1 medium (NCMA, Bigelow Laboratory for Ocean Sciences, USA). The enrichment culture was purified by dilution to 10 cells per well in a 96 deep-well plate (Eppendorf) and incubated under white light (100 μE m^−2^ s^−1^) in a 12:12 h light: dark cycle at 4°C. Cell growth was observed by the development of coloration after a few weeks. Culture purity was assessed by flow cytometry (Becton Dickinson, Accuri C6). After confirmation of the purity, the culture was transferred in a 50 ml ventilated flask (Sarstedt). Cultures are maintained in the Roscoff Culture Collection (http://roscoff-culture-collection.org/) in K/2 (Keller et al. 1987) or L1 medium at 4°C under a 12:12 h light: dark cycle at 100 μE light intensity. RCC2285 has been lost from culture since molecular analyses (described below) were performed. For pigment analysis and electron microscopy, RCC2288 was grown at 7°C under continuous light at 100 μE intensity in L1 medium prepared using autoclaved seawater from offshore Mediterranean Sea water diluted 10% with MilliQ water and filtered prior to use through 0.22 μm filters.

### Sequences

Nuclear 18S rRNA and the Internal Transcribed Spacers (ITS) 1 and 2, as well as the 5.8S rRNA gene were retrieved from GenBank for strains RCC2288, RCC2497 and RCC2285 (Balzano et al. 2012a). For RCC5418 and RCC5150 (*M. antarctica*), cells were harvested in exponential growth phase and concentrated by centrifugation. Total nucleic acids were extracted using the Nucleospin Plant II kit (Macherey-Nagel, Düren, DE) following the manufacturer’s instructions. The nearly full length nuclear 18S rRNA gene (only RCC5418) and the region containing the Internal Transcribed Spacers (ITS) 1 and 2, as well as the 5.8S rRNA gene were obtained by PCR amplification using universal primers (Supplementary Table 1).

PCR products were directly sequenced at the Macrogen Company (Korea) and sequences have been deposited to Genbank under accession numbers MH516003, MH516002 and MH542162.

### Phylogenetic analyses

Nuclear 18S rRNA sequences belonging to members of Mamiellophyceae were retrieved from GenBank (http://www.ncbi.nlm.nih.gov/). Two environmental sequences (similar to strain sequences) were included in addition to the sequences obtained from the cultures. Sequences were also obtained for the ITS2 region located between the 5S and 23S rRNA genes. However, no environmental sequences were available to be included in the 18S/ITS phylogenetic analyses.

Nuclear 18S rRNA and ITS2 sequences were aligned with MAFFT using the E-INS-i and G-INS-i algorithms respectively (Katoh and Toh 2008). For each sequence within the ITS2 alignment, the preliminary secondary structure annotated in dot-bracket format was associated, generating a Vienna file, which was imported to 4SALE (Seibel et al. 2008). The final alignment was edited on the basis of conserved secondary structures. The nuclear 18S rRNA and ITS2 sequences from the Mamiellaceae members were concatenated using Geneious 10.2.5 (Kearse et al. 2012).

Phylogenetic reconstructions with two different methods, maximum likelihood (ML) and Bayesian analyses, were performed using the nuclear Mamiellophyceae 18S rRNA and Mamiellaceae concatenated 18S/ITS2 alignments. The K2 + G + I model was selected for both sequence datasets based on the substitution model selected through the Akaike information criterion (AIC) and the Bayesian information criterion (BIC) options implemented in MEGA 6.06 (Tamura et al. 2013). ML analysis was performed using PhyML 3.0 (Guindon et al. 2010) with SPR (Subtree Pruning and Regrafting) tree topology search operations and approximate likelihood ratio test with Shimodaira-Hasegawa-like procedure. Markov chain Monte Carlo iterations were conducted for 1,000,000 generations sampling every 100 generations with burning length 100,000 using MrBayes 3.2.2 (Ronquist and Huelsenbeck 2003) as implemented in Geneious (Kearse et al. 2012). Nodes were considered as well supported when SH-like support values and Bayesian posterior probabilities were higher than 0.8 and 0.95 respectively. The same criteria were used to represent the sequences on the phylogenetic trees. Alignments are available at https://doi.org/10.6084/m9.figshare.7472153.v1.

### ITS2 secondary structure

ITS2 secondary structure from the strains listed in Table 1 were predicted using the Mfold web interface (Zuker 2003) under the default options with the folding temperature fixed at 37°C, resulting in multiple alternative folding patterns per sequence. The preliminary structure for each sequence was chosen based on similarities found among the other structures proposed for Mamiellophyceae (Marin and Melkonian 2010, Simon et al. 2017) as well as on the presence of previously defined ITS2 hallmarks defined by Coleman (Mai and Coleman 1997, Coleman 2000, 2003, 2007). Exported secondary structures in Vienna format and the respective nucleotide sequences were aligned, visualized using 4SALE version 1.7 (Seibel et al. 2008) and manually edited through extensive comparative analysis of each position (nucleotide) in sequences from representatives of the Mamiellophyceae. The ITS2 synapomorphy analysis was confined to those positions, which formed conserved base pairs in all members of the Mamiellaceae order and the resulting intramolecular folding pattern (secondary structure) of *Mantoniella* was drawn using CorelDRAW X7. A Vienna file containing the ITS2 sequences and secondary structure is available at https://doi.org/10.6084/m9.figshare.7472153.v1.

**Table 1.** Strains used in this study. RCC: Roscoff Culture Collection (www.roscoff-culture-collection.org). 18S rRNA and ITS show Genbank accession numbers. Strains in bold used to describe the new species.

| Strain ID | Strain name | Oceanic Region | Latitude | Longitude | Depth of isolation (m) | 18S rRNA | ITS | Remark |
| --- | --- | --- | --- | --- | --- | --- | --- | --- |
| RCC2285 | MALINA E43.N1 | Beaufort Sea | 70° 34' N | 145° 24' W | 0 | JF794053 | JQ413368 | Strain lost |
| RCC2288 | MALINA E47.P2 | Beaufort Sea | 70° 30' N | 135° 30' W | 0 | JN934679 | JQ413369 |  |
| RCC2497 | MALINA E47.P1 | Beaufort Sea | 70° 30' N | 135° 30' W | 0 | KT860921 | JQ413370 |  |
| RCC5418 | GE_IP_IC_DIL_490 | Baffin Bay | 67° 28' N | 63° 46' W | surface ice | MH51600<br>3 | MH54216<br>2 |  |

### Screening of environmental 18S rRNA sequencing datasets

High-throughput sequencing metabarcodes (V4 and V9 hypervariable regions) were obtained from several published polar studies, as well as from the global sampling efforts Tara Oceans and Ocean Sampling Day (OSD) (see Supplementary Table 2 for the full details and references for each project). We screened these data as well as GenBank by BLASTn (98% identity cut-off) using RCC2288 18S rRNA gene sequence as the search query. We aligned the retrieved environmental sequences and metabarcodes with that of RCC2285, RCC2288, RCC2497, and RCC5418 using MAFFT as implemented in Geneious version 10.0.7 (Kearse et al. 2012). This allowed the determination of sequence signatures diagnostic of this species for both V4 and V9 (Supplementary Figures 1 and 2). The oceanic distribution of stations where cultures, clones and metabarcodes having these signatures, as well as the stations from the metabarcoding surveys where no matching metabarcodes have been found, were plotted with the R libraries ggplot2 and rworldmap. The R script is available at https://vaulot.github.io/papers/RCC2288.html.

### Light microscopy

Cells were observed using an Olympus BX51 microscope (Olympus, Hamburg, Germany) with a 100× objective using differential interference contrast (DIC) and imaged with a SPOT RT-slider digital camera (Diagnostics Instruments, Sterling Heights, MI, USA).

For video-microscopy, cultures from RCC2288 and RCC2497 were observed with an inverted Olympus IX70 inverted microscope using an x40 objective and equipped with an Infinity X camera (https://www.lumenera.com/products/microscopy/infinityx-32.html). Short sequences were recorded and edited with the Video de Luxe software (http://www.magix.com/fr/video-deluxe/). Films were uploaded to Youtube (https://www.youtube.com/channel/UCsYoz-aSJlJesyDNj6ZVolQ/videos). The recording protocol is available at https://www.protocols.io/private/1fbc54800109d5f44e88574f40194ed1

### Transmission Electron Microscopy

Positively stained whole mount cells were prepared described (Moestrup 1984) where cultures were directly deposited on formvar coated copper grids and stained with 2% uranyl acetate. TEM thin-sections was performed as previously described (Derelle et al. 2008). Briefly, fixed RCC2288 cells (1% glutaraldehyde) from an exponentially growing culture were suspended in molten (37°C) 1% low melting point agarose. The agarose cell plug was fixed, washed, dehydrated in ethanol and embedded in Epon 812. Ultra-thin sections (80–90 nm) were placed on a 300 mesh copper grid and stained with uranyl acetate for 15 min, followed by lead citrate staining for 2 min. The cells were visualised with Hitachi H 7500 and H-9500 transmission electron microscopes.

### Pigment analysis

Pigments were extracted from RCC2288 cells in late exponential phase as previously described (Ras et al. 2008). Briefly, cells were collected on 0.7 μm particle retention size filters (GF/F Whatman), pigments extracted for 2 hours in 100% methanol, then subjected to ultrasonic disruption and clarified by filtration through 0.2 μm pore-size filters (PTFE). Pigments were detected using high performance liquid chromatography (HPLC, Agilent Technologies 1200) over the 24 h after the extraction.

## Results and Discussion

### Taxonomy section

#### *Mantoniella beaufortii* Yau, Lopes dos Santos and Eikrem sp. nov.

Diagnosis: Cells round measuring 3.7 ± 0.4 μm in diameter with one long (16.3 ± 2.6 μm) and one short reduced flagellum (~1 μm). Cell body and flagella covered in imbricated spiderweb scales. Flagellar hair scales present composed of two parallel rows of subunits. Long flagellum tip has tuft of three hair scales. Scales produced in Golgi body. Golgi body located beneath and to the side of basal bodies. One green chloroplast with pyrenoid surrounded by starch and a stigma composed of a single layer of oil droplets (ca 0.1 μm). Ejectosomes composed of fibrils located at periphery of cell. Cell bodies with sub-quadrangular to oval scales (~0.2 μm). Body scales heptaradial, with seven major spokes radiating from center, number of spokes increasing in number towards the periphery. Six or more concentric ribs divide the scale into segments. Flagella with hexaradial oval scales composed of six spokes increasing in number towards the periphery. Six or more concentric ribs divide the scale into segments. Combined nucleotide sequences of the 18S rRNA (JN934679) and rRNA ITS2 (JQ413369) are species specific. In ITS2 of the nuclear encoded rRNA operon, base pair 4 of Helix I is C-G instead of U-A and base pair 22 of Helix IV is G-C instead of G-U.

##### Holotype

Plastic embedding deposited at the Natural History Museum, University of Oslo, accession numbers O-A-10010. Figure 4 shows cells from the embedding. Culture deposited in The Roscoff Culture Collection as RCC2288.

##### Type locality

Strain RCC2288 was isolated from surface water sampled from the Beaufort Sea in the Arctic Ocean (70°30’N, 135°30’W) on 14th July 2009.

##### Etymology

Named for its geographical provenance.

#### *Mantoniella baffinensis* Yau, Lopes dos Santos and Eikrem sp. nov.

Diagnosis: Cells measuring 4.7 ± 0.5 μm with a long flagellum of 21.8 ± 5.1 μm and one short reduced flagellum (~1 μm). Cell body and flagella covered in imbricated spiderweb scales. Flagellar hair scales present composed of two parallel rows of subunits. Long flagellum tip has tuft of three hair scales. Cell bodies with sub-quadrangular to oval scales (~0.2 μm). Body scales octaradial with eight major radial spokes radiating from center, number of spokes increasing in number towards the periphery. Seven or more concentric ribs divide the scale into segments. Flagella with heptaradial, oval scales composed of seven spokes increasing in number towards the periphery. Six or more concentric ribs divide the scale into segments. Combined nucleotide sequences of the nuclear 18S rRNA (MH516003) and rRNA ITS2 (MH542162) are species specific. In ITS2 of the nuclear encoded rRNA operon, base pair 9 of Helix I is C-G instead of A-U, base pair 15 of Helix II is G-C instead of A-U and base pair 22 of Helix IV is C-G instead of G-U.

##### Holotype

Plastic embedding deposited at the Natural History Museum, University of Oslo, accession number O-A-10011. Culture deposited in The Roscoff Culture Collection as RCC5418.

##### Type locality

Strain RCC5418 was isolated from surface sea ice sampled off the coast of Baffin Island, in the Baffin Bay (67°28’N, 63°46’W) on 23th May 2016.

##### Etymology

Named for its geographical provenance.

### Phylogeny and ITS signatures

The phylogenetic tree based on nearly full-length nuclear 18S rRNA sequences obtained from the novel polar strains RCC2288, RCC2285, RCC2497 and RCC5418 (Table 1), and environmental sequences retrieved from GenBank indicate that these strains belong to the family Mamiellaceae (Supplementary Figure 3). The analysis also recovers the major genera within the order Mamiellales: *Bathycoccus, Ostreococcus, Micromonas, Mantoniella* and *Mamiella* (Marin and Melkonian 2010). Dolichomastigales and Monomastigales were the basal orders in the class Mamiellophyceae with the *Monomastix opisthostigma* type species used as an outgroup. The three strains isolated during the MALINA cruise in the Beaufort Sea (RCC2485, RCC2288 and RCC2497) and the strain from the Baffin Bay (RCC5418) form a well-supported clade together with two environmental sequences (clone MALINA St320 3m Nano ES069 D8 and clone 4-E5), also originated from Arctic Ocean samples. The two described *Mantoniella* species (*M. squamata* and *M. antarctica*) are not monophyletic in our analysis using the nuclear 18S rRNA, as observed by Marin and Melkonian (2010) (Supplementary Figure 3).

In contrast, the phylogenetic tree based on concatenated 18S/ITS2 alignments suggests that our strains belong to the genus *Mantoniella* (Figure 1). The topology of the concatenated 18S/ITS tree, in addition to grouping our strains within *Mantionella*, is consistent with a recent nuclear multigene phylogeny based on 127 concatenated genes from related Chlorophyta species that included RCC2288 (Lopes dos Santos et al. 2017b). This indicates the 18S/ITS2 tree reflects the evolutionary history of the nuclear genome supporting the position of *Mantoniella* and our strains diverging from the same common ancestor.

**Figure 1.**
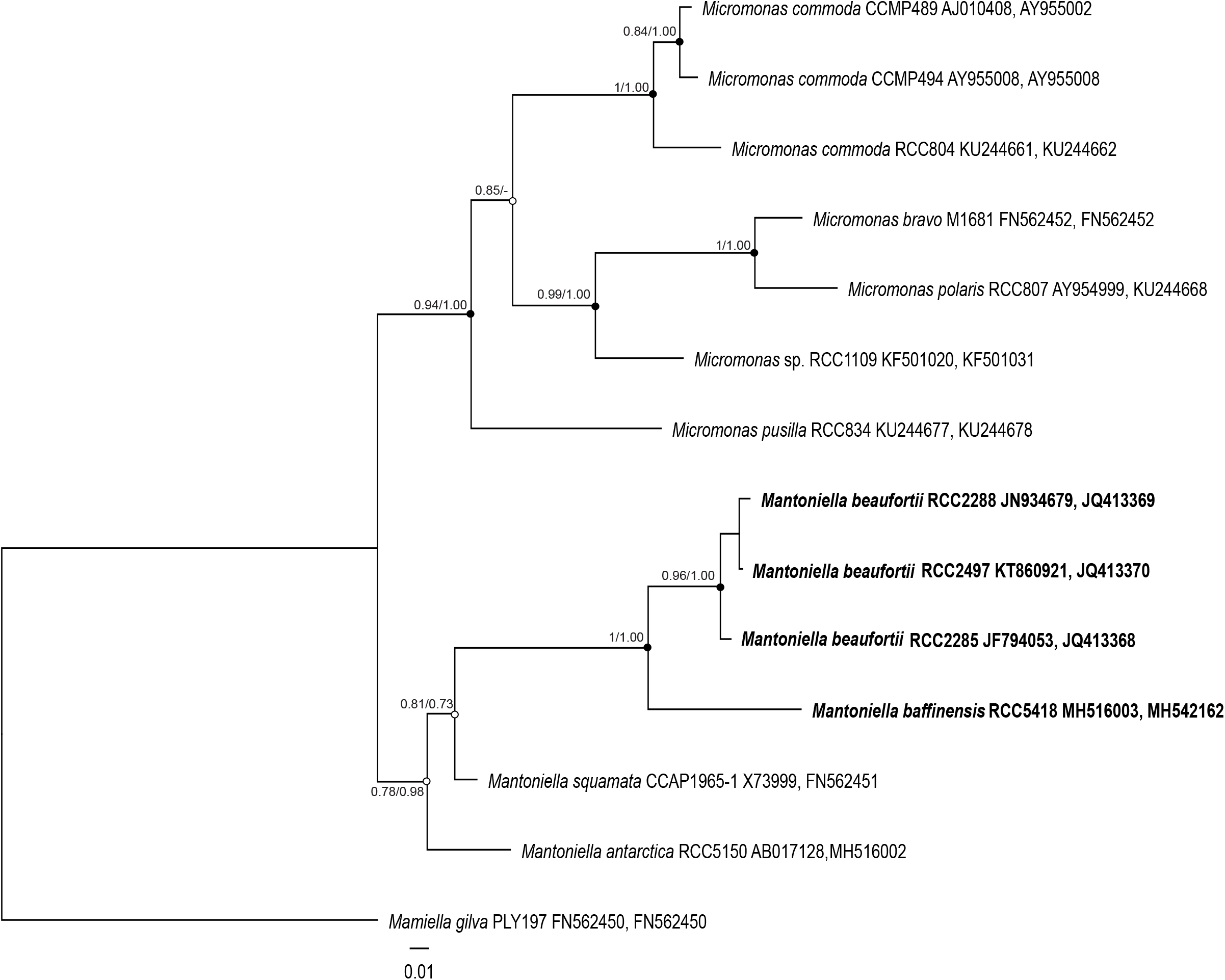
Maximum-likelihood tree inferred from concatenated 18S/ITS2 sequences of Mamiellaceae. Solid dots correspond to nodes with significant support (> 0.8) for ML analysis and Bayesian analysis (> 0.95). Empty dots correspond to nodes with non-significant support for either ML or Bayesian analysis, or both.

The average distance between strains RCC2485, RCC2288 and RCC2497 is low (0.5% of segregating sites over the near full-length 18S rRNA gene), suggesting that these strains correspond to a single species that we name *Mantoniella beaufortii* (see taxonomy section). In contrast, the well-supported placement of strain RCC5418 on an earlier diverging branch within the *Mantoniella* clade, as well as the 1% average distance between RCC5418 and the other strains, suggests it represents another species, named here *Mantoniella baffinensis*.

To substantiate the description of *M. beaufortii* and *M. baffinensis* as new species, we investigated ITS2 synapomorphies of the different *Mantoniella* species. Several studies have shown the power of using ITS2 sequences in delimiting biological species, especially in microalgae studies (e.g. Coleman 2007, Caisová et al. 2011, Lopes dos Santos et al. 2017b, Simon et al. 2017). For example, ITS sequencing contributed to distinguishing the Arctic diatom *Chaetoceros neogracilis* from an Antarctic *Chaetoceros* sp. that share nearly identical 18S rRNA genes (Balzano et al. 2017). Moreover, nucleotide diversity within the ITS2 allowed the identification of four distinct populations within the Beaufort Sea. The computed ITS2 secondary structure of the new *Mantoniella* strains contains the four helix domains found in many eukaryotic taxa (Supplementary Figure 4), in addition to the presence of Helix B9. The intramolecular folding pattern of the ITS2 transcript from *M. beaufortii* and *M. baffinensis* is very similar to the one from *M. squamata* and *M. antarctica* (Supplementary Figure 4). The universal hallmarks proposed by Mai and Coleman et al. (1997) and Schultz et al. (2005) are present in Helices II and III of the Mamiellaceae. These are the Y-Y (pyrimidine-pyrimidine) mismatch at conserved base pair 7 in Helix II (Figure 2) and YRRY (pyrimidine-purine-pyrimidine) motif at conserved positions 28–31 on the 5’ side of Helix III (Supplementary Figure 5A). In all four strains, the Y-Y mismatch is represented by the pair U-U and the YRRY motif by the sequence UGGU.

**Figure 2.**
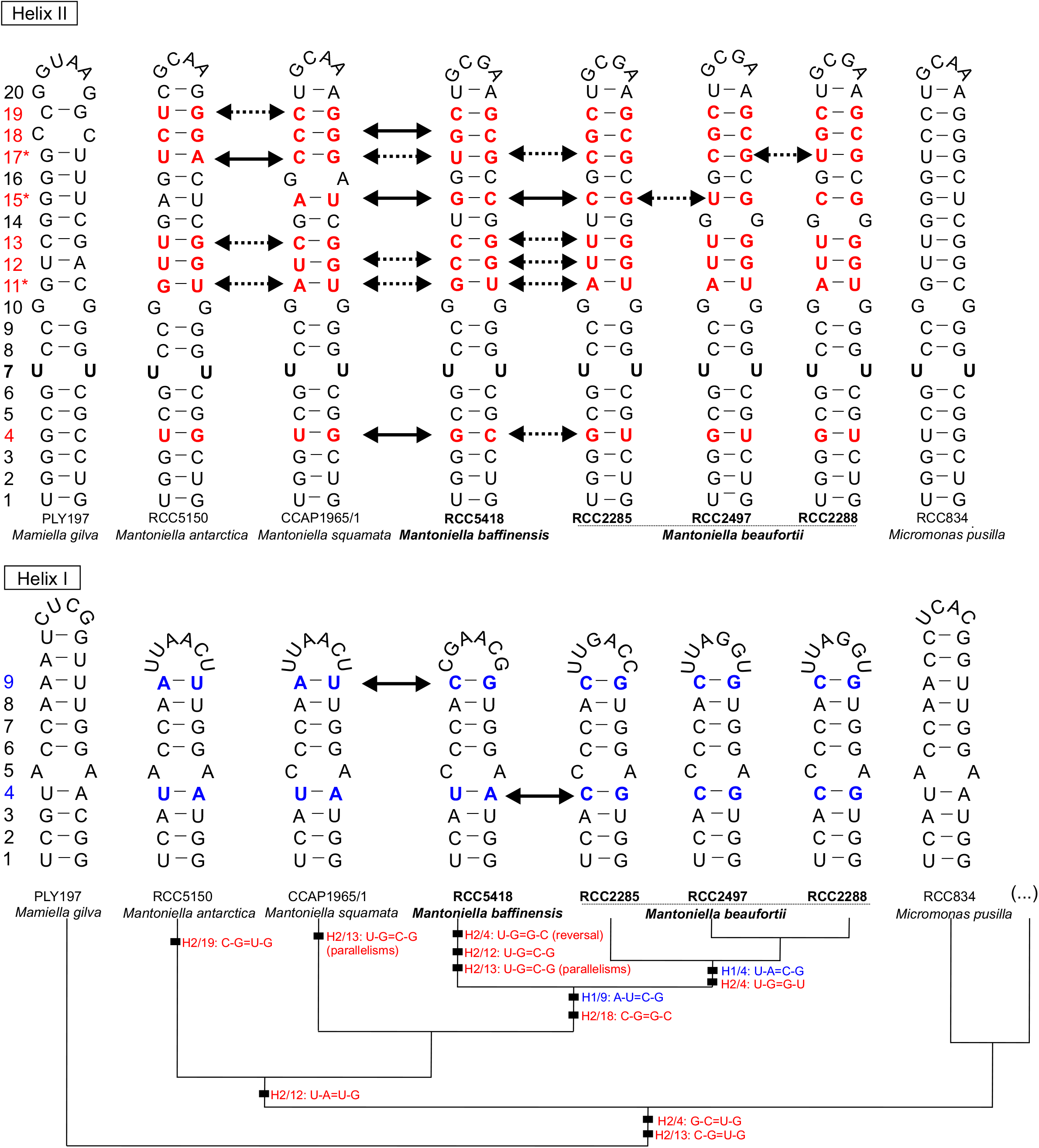
Molecular signatures of *Mantoniella* species based on comparison of ITS2 secondary structures within Mamiellaceae. Signatures in Helix I are shown in blue and Helix II in red. The conserved base pairs among the different groups are numbered. Compensatory base changes (CBCs) and hemi-CBCs (hCBSs) are highlighted by solid and dotted arrows respectively. Hypervariable positions are marked by an asterisk (*). Ellipsis (…) represent the other clades and species of *Micromonas*. The pyrimidine-pyrimidine (Y-Y) mismatch in Helix II is shown in bold black. Single nucleotide substitutions are shown by grey nucleotides. Identified homoplasious changes are shown as parallelisms and reversals.

The structural comparison at each base pair position within the ITS2 helices identified several compensatory base changes (CBCs) and single-side changes or hemi-CBCs (hCBCs), as well as conserved base pair positions among *Mantoniella* species (Supplementary Figure 4). Note that we only considered hCBCs at positions where the nucleotide bond was preserved. No CBCs was found between the three *M. beaufortii* strains consistent with their designation as a single species. However, three hCBCs were detected in Helix II at positions 15 and 17 (Figure 2) and Helix III at position 12 (Supplementary Figure 5A). Three CBCs were detected in Helices I (position 4), II (position 15) and IV (position 22) between *M. beaufortii* and *M. baffinensis*, supporting the separation of these strains into two distinct species (Figure 2 and Supplementary Figure 4). When possible, the evolutionary steps of the identified CBCs and hCBCs were mapped upon branches of the Mamiellaceae phylogenetic tree that was constructed based on the concatenated 18S/ITS2 (Figure 2 and Supplementary Figure 4) to distinguish synapomorphies from homoplasious changes (e.g. parallelisms and reversals). Few hypervariable positions showing several changes (CBCs and hCBCs) could not be unambiguously mapped upon the tree.

### Morphology and ultrastructure

Under light microscopy, the cells of the new strains are green and round with one long and one short reduced flagellum (~1 μm), which are inserted almost perpendicularly to the cell (Figure 3). They swim with their flagella directed posteriorly, pushing the cell. Occasionally the cells cease movement, pirouette and take off again in a different direction (Supplementary Material 1–3). All strains possess a stigma, visible in light microscopy as a red eyespot located opposite the flagella. Although there are no morphological characters that are unique to the mamiellophyceans and shared by all of its members, the new strains closely resemble *Mantoniella* and *Mamiella*, which are similarly small round bi-flagellated cells (see Supplementary Table 3 for morphological characters in described Mamiellophyceae). However, the flagella of *Mamiella* are of equal or near equal lengths (Moestrup 1984), so clearly the unequal flagella observed in our strains conform with described *Mantoniella* species, *M. squamata* and *M. antarctica* (Barlow and Cattolico 1980, Marchant et al. 1989). The new strains are thus morphologically indistinguishable by light microscopy from *Mantoniella* species, supporting their placement in the genus.

**Figure 3.**
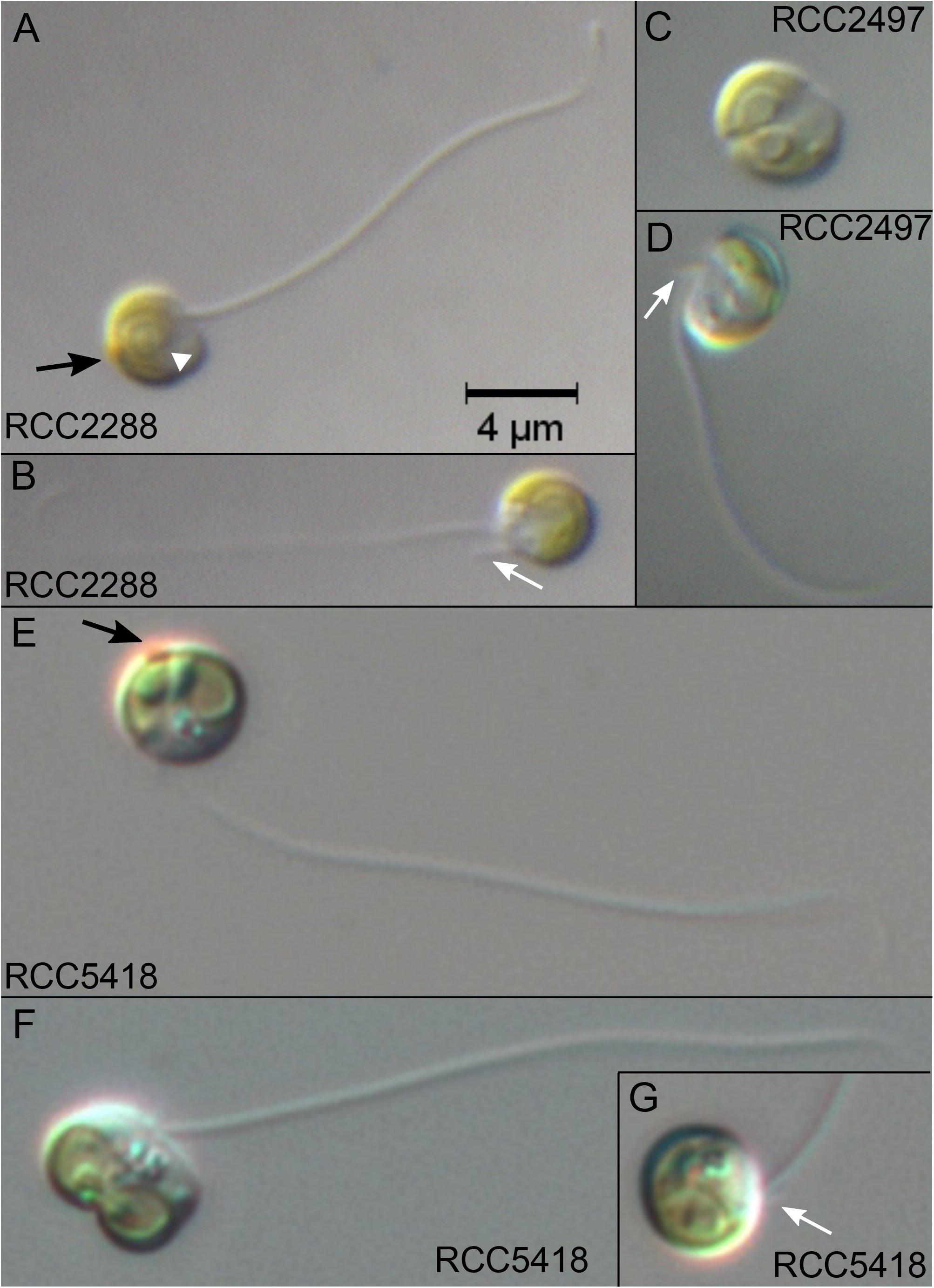
Light microscopy images of the new *Mantoniella* strains. All strains have round cell morphology, visible red stigma (black arrow), a long and short flagellum (white arrow) and one chloroplast with a pyrenoid (white arrowhead). Scale bar is 4 μm for all images. **(A-B)** *M. beaufortii* RCC2288. **(C-D)** *M. beaufortii* RCC2497 during cell division and single cell showing long and short flagellum. **(E-G)** *M. baffinensis* RCC5418 single cell **(E)**, during cell division **(F)** and cell showing the short flagellum (**G** inset).

The new strains are in the size range (Table 2) reported for *M. squamata* (3–6.5 μm) and *M. antarctica* (2.8–5 μm) (Manton and Parke 1960, Marchant et al. 1989). Nonetheless, *M. beaufortii* strains are significantly smaller than *M. baffinensis* in cell diameter and average long flagellum length (Table 2) providing a means to distinguish the two new *Mantoniella* species from each other with light microscopy.

**Table 2.** Cell diameter and long flagellum lengths measured for *M. beaufortii* (RCC2288 and RCC2497) and *M. baffinensis* (RCC5418). n = number of cells measured and SD = standard deviation.

| Strain | min | max | mean | median | stdev | n |
| --- | --- | --- | --- | --- | --- | --- |
| **Cell diameter (µm)** |  |  |  |  |  |  |
| RCC2288 | 2.89 | 4.98 | 3.77 | 3.70 | 0.41 | 60 |
| RCC2497 | 3.15 | 4.74 | 3.87 | 3.77 | 0.39 | 39 |
| RCC5418 | 3.54 | 5.69 | 4.66 | 4.66 | 0.51 | 69 |
| **Long flagellum length (µm)** |  |  |  |  |  |  |
| RCC2288 | 12.93 | 21.47 | 16.27 | 15.99 | 2.63 | 11 |
| RCC2497 | 11.91 | 21.25 | 16.31 | 17.07 | 2.71 | 12 |
| RCC5418 | 11.27 | 32.59 | 21.78 | 21.29 | 5.14 | 25 |

Transmission Electron Microscopy (TEM) of thin sections (Figure 4) and whole mounts (Figure 5) of the new strains provided details of their internal and external morphological features. The single chloroplast is green and cup-shaped with a pyrenoid surrounded by starch tubules running through the pyrenoid. The stigma is composed of a single layer of oil droplets (approximately 0.1 μm in diameter) (Figure 4A) and located at the periphery of the chloroplast facing the cell membrane, conforming to the description of the family Mamiellaceae (Marin and Melkonian 2010). Several large ejectosomes composed of fibrils are present at the cell periphery (Figure 4D and E). They are common in the Mamiellales (Moestrup 1984, Marchant et al. 1989) and are perhaps used to deter grazers.

**Figure 4.**
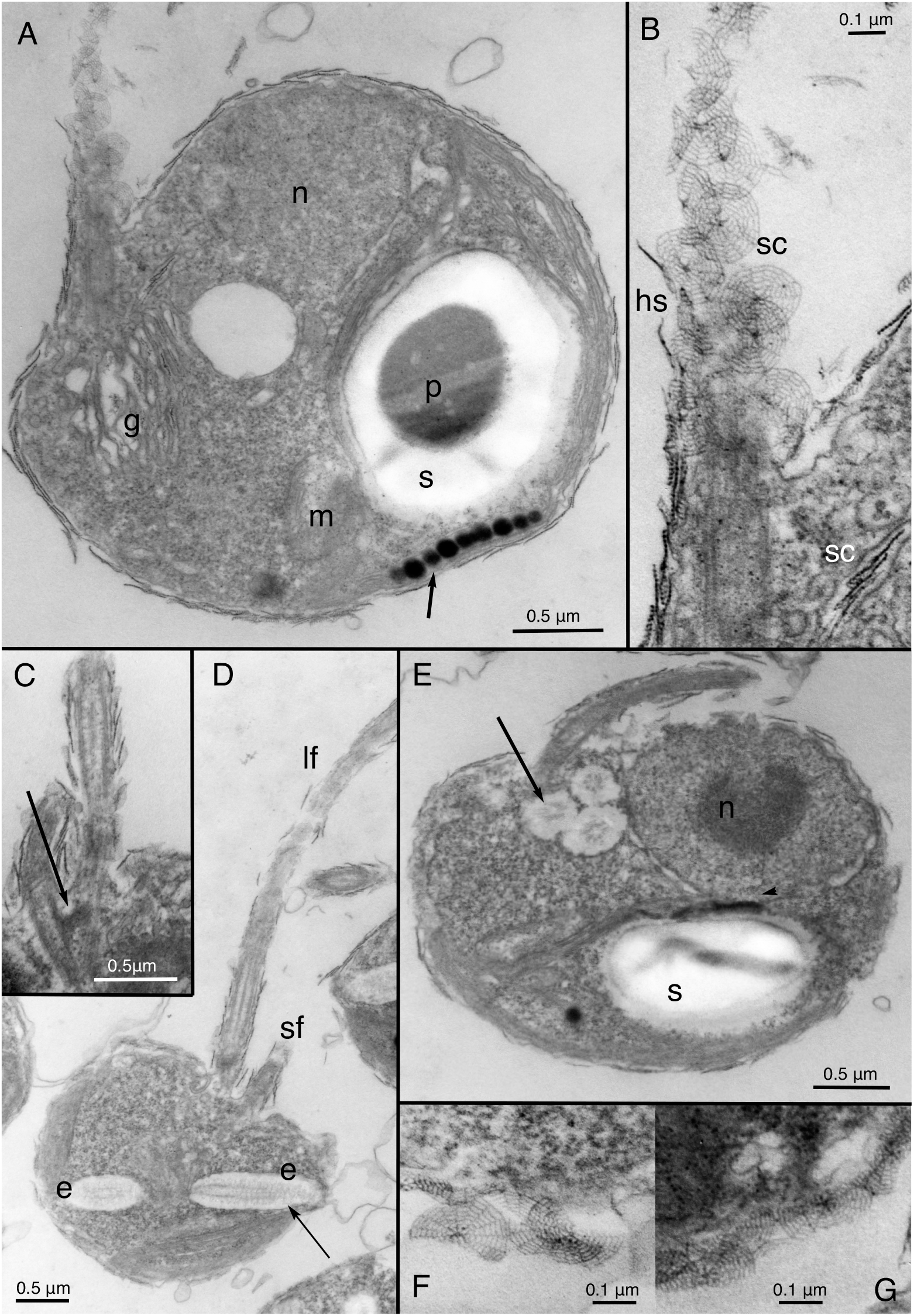
TEM thin sections of *M. beaufortii* RCC2288. (**A**) Internal cell structure showing organelles and stigma (black arrow). (**B**) Detail of the hair and spiderweb scales covering the long flagellum. Scales are produced in the Golgi body. (**C**) Detail of the flagellar base (black arrow). (**D**) Cell with long and short flagella and longitudinal section of the ejectosomes (black arrow). (**E**) Cross section of ejectosomes (black arrow). (**F**) and (**G**) Body scales made up of radiating and concentric ribs. Abbreviations: e=ejectosome, g=Golgi, s=starch granule, m=mitochondrion, n=nucleus, p=pyrenoid, hs=hair scale, sc=scale, lf=long flagellum and sf=short flagellum.

**Figure 5.**
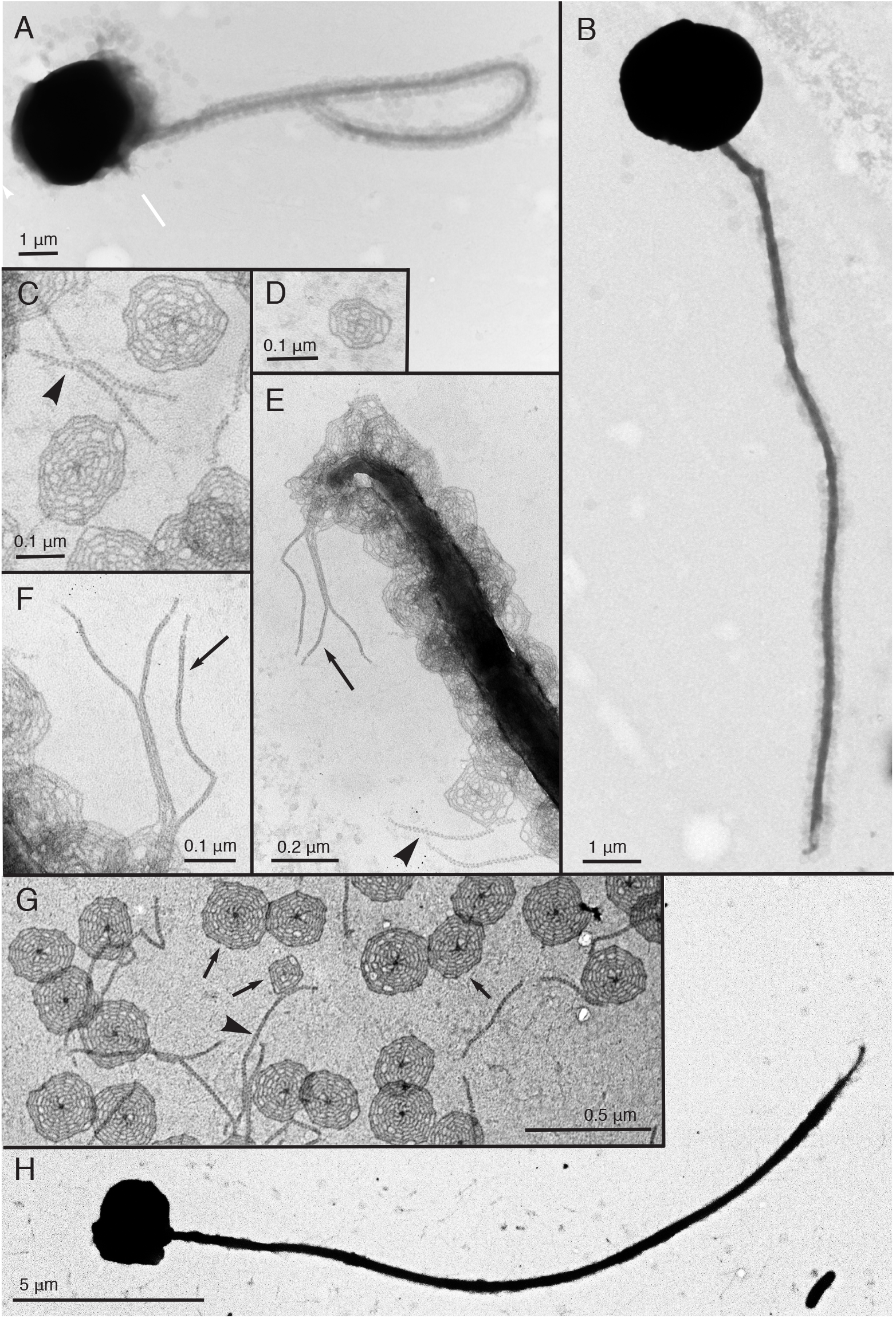
Transmission electron micrographs of whole-mounts of the new *Mantoniella* strains. **(A-E)** *M. beaufortii*. (**A**) Whole cells of strain RCC2288, indicating the short flagellum (white arrow), and (**B**) RCC2497. (**C**) Detached flagellar spiderweb-like scales and hair scales (black arrowhead). (**D**) Detail of small tetraradial body scale. (**E**) Imbricated scales and hair scales covering the long flagellum. A tuft of three hair scales on the tip of the long flagellum (black arrow) (**F**) Detail of the tuft of hair scales (black arrow). **(G-H)** *M. baffinensis* RCC5418. (**G**) Small and large body scales (black arrows) and flagellar hair scales (black arrowhead) and (**H**) whole cell.

One of the most salient features of the Mamiellophyceae is the presence of organic scales covering the cell, the most common of which comprise radiating and concentric ribs resembling spiderwebs that are present in the scale-bearing Mamiellales (*Bathycoccus*, *Mamiella* and *Mantoniella*), as well as *Dolichomastix* (Supplementary Table 3). We examined the whole mounts of the new *Mantoniella* species to establish the presence of scales and determine if they were morphologically distinguishable from related species, as *M. antarctica* (Marchant et al. 1989) and *M. gilva* (Moestrup 1984) each have a unique type that differentiate them from other Mamiellales.

The flagella and cell bodies of the new strains are covered in imbricated spiderweb-like scales (Figure 5) measuring approximately 0.2 μm. The scales are produced in the Golgi body (Figure 4B). The body scales are sub-quadrangular to oval whereas the flagellar scales are oval (Figure 5). Spiderweb scales have 6–8 major spokes radiating from the center with the number of spokes increasing in number towards the periphery and six or more concentric ribs dividing the scale into segments. In addition, there are some small scales (approximately 0.1 μm) on the cell body composed of four spokes (increasing to eight) and separated by four, more or less concentric, ribs (Figure 5D, G). The flagella are also covered by lateral hair scales, which are composed of two parallel rows of globular subunits. At the tip of the long flagellum there is a tuft of three hair scales, for which the subunits are more closely packed together than the lateral hair scales (Figure 5). The hair scales of the new strains are identical to the “Tetraselmis-type” T-hairs previously described in *Mantoniella* and *Mamiella* (Marin and Melkonian 1994). This structure is otherwise only seen in *Dolichomastix lepidota* and differs from the smooth tubular T-hairs of *Dolichomastix tenuilepis* and *Crustomastix* (Marin and Melkonian 1994, Zingone et al. 2002)(Supplementary Table 3).

Comparison of the spiderweb scales between *Mantoniella* species (Table 3) shows the new species differ significantly from *M. antarctica*, which possesses lace-like scales with six or seven radial ribs with very few concentric ribs (Marchant et al. 1989). Morphologically, the spiderweb scales of the new species most resemble *M. squamata*, which has large heptaradial flagellar scales, octaradial body scales and a few additional small tetraradial body scales (Marchant et al. 1989). Indeed, the spiderweb scales of *M. baffinensis* (Figure 5) are structurally indistinguishable from *M. squamata*. In contrast, *M. beaufortii* shares the small tetraradial body scales but possesses hexaradial flagellar scales and heptaradial body scales, potentially allowing it to be differentiated from the other *Mantoniella* based on the number of radial spokes of the spiderweb scales.

**Table 3.** Comparison of *Mantoniella* spp. scale types.

| Species | Flagellar scales | Body scales |
| --- | --- | --- |
| *Mantoniella squamata* | spiderweb-like heptaradial | spiderweb-like large octaradial and small rare tetraradial |
| *Mantoniella antarctica* | lace-like heptaradial | lace-like hexaradial and smaller heptaradial |
| *Mantoniella beaufortii* | spiderweb-like hexaradial | spiderweb-like large heptaradial and small rare tetraradial |
| *Mantoniella baffinensis* | spiderweb-like heptaradial | spiderweb-like large octaradial and small rare tetraradial |

### Pigment composition

Pigment to chlorophyll *a* ratios of *M. beaufortii* RCC2288 were compared to a selection of other Chlorophyta species (Figure 6, Supplementary Table 4) from previous studies (Latasa et al. 2004, Lopes dos Santos et al. 2016), as pigments are useful phenotypic traits. Chlorophyll *a* and b, characteristics of Chlorophyta, were detected, as well as the basic set of carotenoids found in the prasinophytes: neoxanthin, violaxanthin, lutein, zeaxanthin, antheraxanthin and β-carotene. The additional presence of prasinoxanthin, micromonal and uriolide places RCC2288 in the PRASINO-3B group of prasinophyte green algae, *sensu* Jeffrey et al. (2011). This pigment-based grouping shows good agreement with the molecular phylogeny of Mamiellales, where the presence of prasinoxanthin, micromonal and the Unidentified M1 pigment are diagnostic of the order (Marin and Melkonian 2010). We did not detect Unidentified M1 in RCC2288, but as our analysis method differed from previous work (Latasa et al. 2004) and we relied on matching its chromatographic and spectral characteristics, its absence requires further confirmation. Notwithstanding, the pigment complement of RCC2288 is identical to other described Mamiellales (Figure 6, Supplementary Table 4), coherent with its classification within this order.

**Figure 6.**
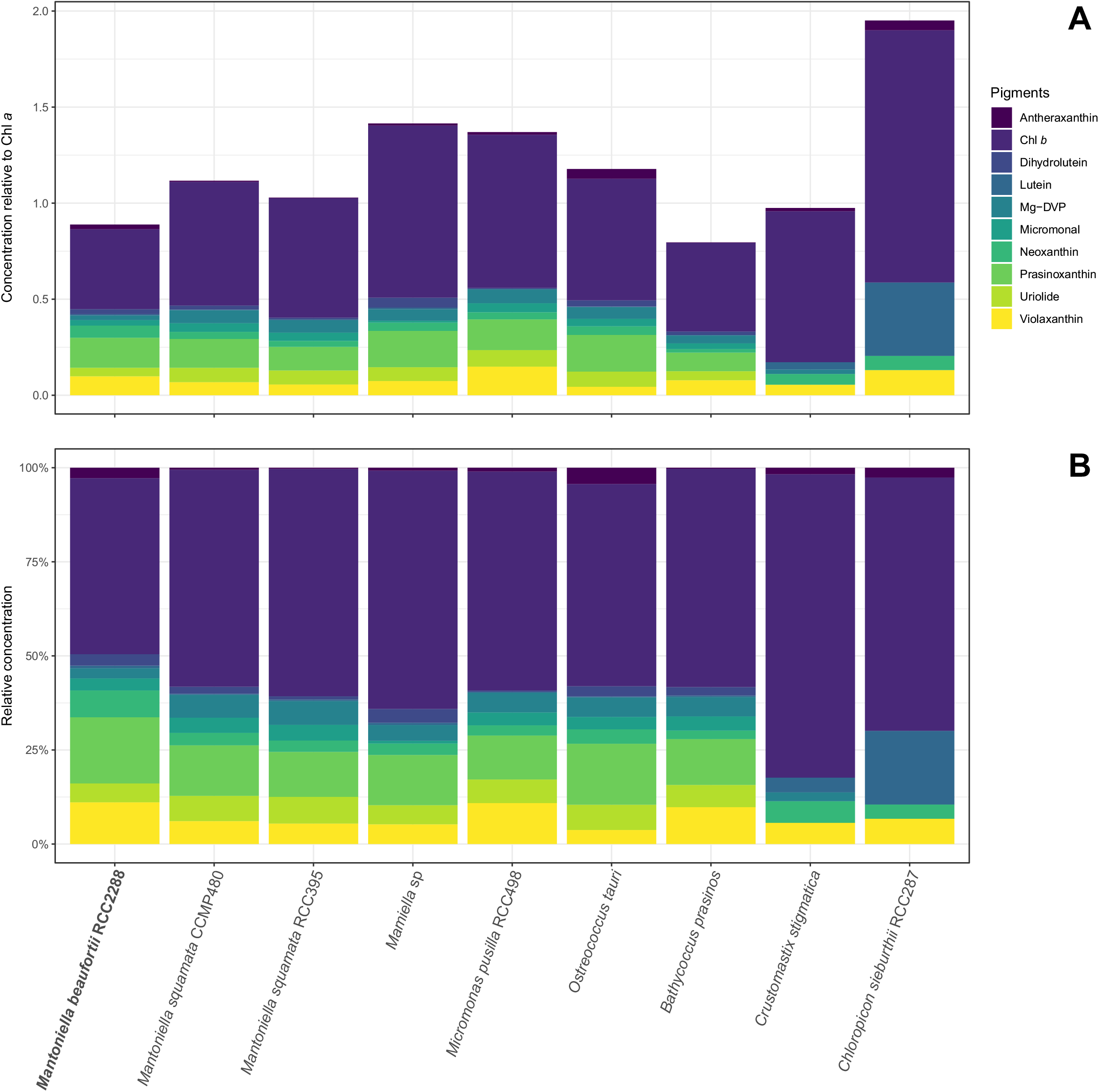
Pigment to chlorophyll *a* ratios in *M. beaufortii* RCC2288 (this study) compared to other Mamiellophyceae species (data from Latasa et al. 2004). (**A**) Cumulative pigment to Chlorophyll *a* ratio of Chlorophyll *b* and abundant carotenoids (excluding α- and β-carotene). (**B**) As for A, but showing relative abundances. Mg-DVP: Mg-24-divinyl pheoporphyrin a5 monmethyl ester.

As noted by Latasa et al. (2004), Mamiellales pigment profiles are remarkably comparable (Figure 6), despite strains being cultured under very different conditions. Only a few carotenoids differed substantially (at least two fold) in relative abundance between *M. beaufortii* and the two other *M. squamata* strains analyzed: the concentration of neoxanthin, antheraxanthin and lutein were higher, whereas that of Mg-DVP and uriolide were relatively lower (Figure 6, Supplementary Table 4). Neoxanthin (associated with the light harvesting complex), as well as antheraxanthin and lutein (both involved in photoprotection), have previously been shown to increase significantly in *M. squamata* grown under continuous light compared to alternating light/dark cycles (Böhme et al. 2002). Therefore, the relatively high ratio of these carotenoids measured in *M. beaufortii* is consistent with growth under continuous light used with RCC2288. Uriolide and Mg-DVP have been observed to increase with light intensity in *M. squamata* (Böhme et al. 2002) and *Micromonas pusilla* (Laviale and Neveux 2011), respectively. Although more physiological data are required to interpret their relative decrease in RCC2288, these pigments are probably most responsive to light conditions (intensity and photoperiod).

Two unknown carotenoids were detected in RC2288, the first one having adsorption peaks at 412, 436 and 464 nm, and the second one at 452 nm (Supplementary Table 5). These were relatively minor components comprising 2.7% and 1.5% of total carotenoids, respectively and may represent carotenoids unique to *M. beaufortii*.

### Environmental distribution

In order to obtain information on the distribution of these two new species, we searched by BLAST both environmental GenBank sequences and published 18S V4 and V9 metabarcodes data sets (Supplementary Table 2). This allowed the retrieval of a few 18S rRNA sequences with higher than 98% similarity to the gene of RCC2288. Alignment of these sequences with other Mamiellophyceae sequences revealed diagnostic positions in both the V4 and V9 hypervariable regions permitting *M. beaufortii* and *M. baffinensis* to be distinguished from other Mamiellophyceae, especially other *Mamiella* and *Mantoniella* species (Supplementary Figures 1 and 2). Signatures from the V4 region were clearer than from V9 due to the fact that for some of the strains, the sequences did not extend to the end of the V9 region (Supplementary Figure 2). In the V4 region, three signatures were observed, one common to both species (A in Supplementary Figure 1), while the other two (B and C in Supplementary Figure 1) differed between *M. beaufortii* and *baffinensis*.

No clone library or metabarcode sequences matched exactly *M. baffinensis*. In contrast, three environmental sequences (KT814860, FN690725, JF698785) from clone libraries had signatures similar to the *M. beaufortii* strains, two from Arctic Ocean water (Figure 7), including one obtained during the MALINA cruise, and one from ice originating from the Gulf of Finland. V4 metabarcodes corresponding to *M. beaufortii* were found in the Ocean Sampling Day data set (Kopf et al. 2015) that includes more than 150 coastal samples at a single station off East Greenland as well as in three metabarcoding studies in the Arctic Ocean, one in the Beaufort Sea performed during the MALINA cruise (Monier et al. 2015), one from Arctic sea ice (Stecher et al. 2016) where it was found at three stations and one from the White Sea (Belevich et al. 2017), also in the sea ice (Figure 7). No metabarcode corresponding to these two new species were found in waters from either the Southern Ocean or off Antarctica (Figure 7 and Supplementary Table 2). No metabarcodes from the V9 region corresponding to the two new species were found in the Tara Oceans data set that covered mostly temperate and subtropical oceanic regions (de Vargas et al. 2015). These data suggest that these species are restricted to polar Arctic regions (although we cannot exclude that they may be found in the future in the Antarctic which has been under-sampled until now) and are probably associated to sea ice although they can be present in the sea water, and that *M. beaufortii is* more wide spread than *M. baffinensis*.

**Figure 7.**
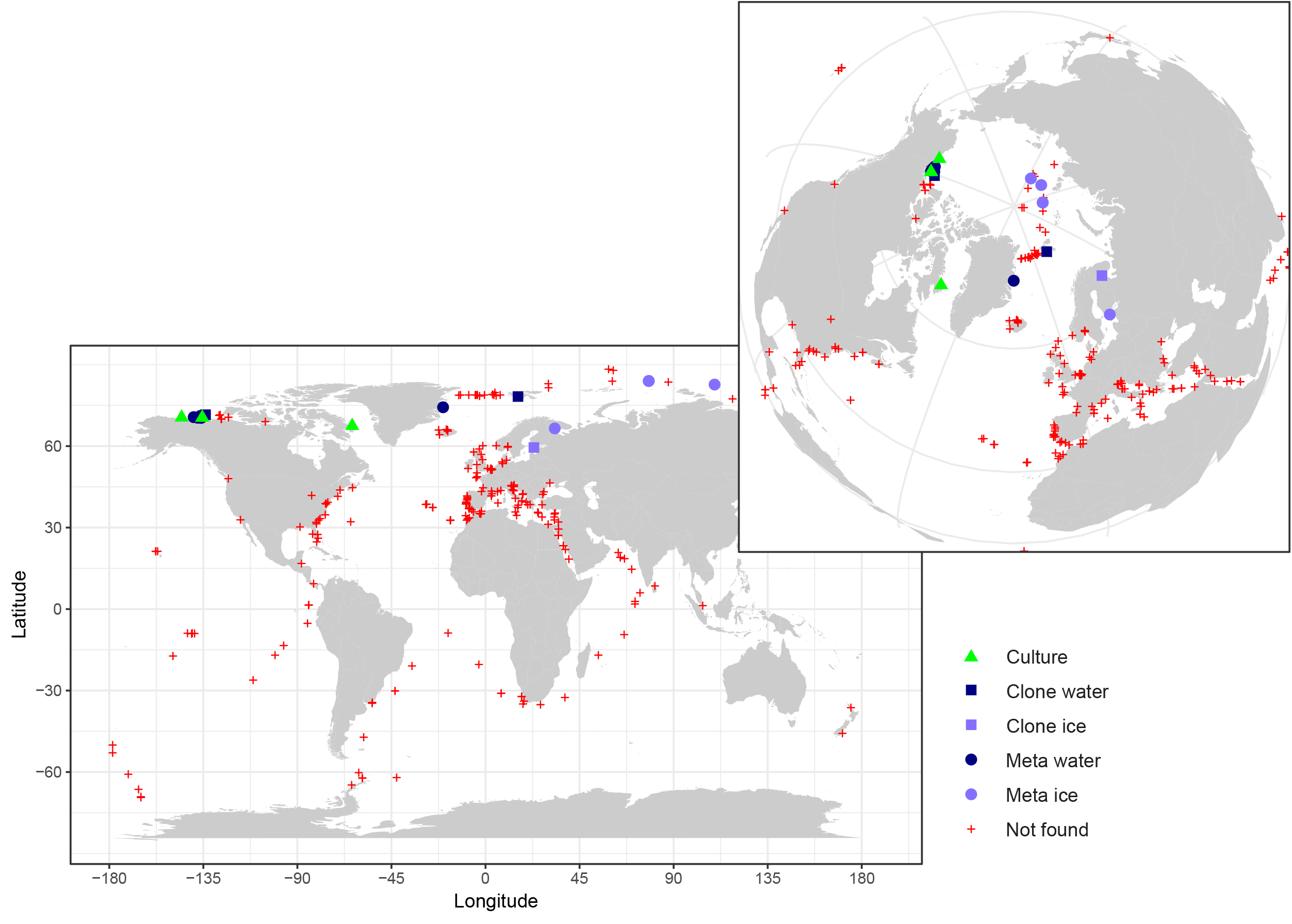
Map of the distribution of *M. beaufortii* in environmental sequence datasets highlighting its prevalence in Arctic samples (inset). The isolation sites of *M. beaufortii* cultures, presence of its 18S rRNA gene sequence in clone libraries (Clone water, Clone ice) and metabarcodes from seawater and ice samples (Meta water, Meta ice) and absence in metabarcodes (Not found) are plotted. For *M. baffinensis*, only its isolation site is indicated in Baffin Bay since no similar environmental sequence was found in the datasets analyzed. Metabarcoding datasets include Ocean Sampling Day, Tara Oceans and polar projects. See Supplementary Table 2 for a full description of the metabarcoding datasets screened.

## Funding

Financial support for this work was provided by the following projects: ANR PhytoPol (ANR-15-CE02-0007) and Green Edge (ANR-14-CE01-0017-03), ArcPhyt (Région Bretagne), TaxMArc (Research Council of Norway, 268286/E40).

## Acknowledgments

We are thankful to Adam Monier, Katja Metfies, Estelle Kilias and Wei Luo for communicating raw metabarcoding data and Sophie Le Panse and Antje Hofgaard for assistance with electron microscopy. We thankful for the support of the BioPIC flow cytometry and microscopy platform of the Oceanological Observatory of Banyuls and of the ABIMS bioinformatics platform at the Roscoff Biological Station.

## Supplementary Figures

Supplementary Figure 1. Alignment of the 18S rRNA gene V4 hypervariable region from *M. beaufortii* and *M. baffinensis* strains (Red and Orange, respectively), environmental clones (Blue) and metabarcodes (Green) with a selection of sequences from closely related Mamiellophyceae. Sequence signatures diagnostic of the two new species are indicated by boxes. The A region is specific of both species while the B and C regions differ between the two species.

Supplementary Figure 2. Alignment of the 18S rRNA gene V9 hypervariable region from *M. beaufortii* and *M. baffinensis* strains (Red and Orange, respectively) and environmental clones (blue) with a selection of closely related Mamiellophyceae sequences. Sequence signatures diagnostic of *M. beaufortii* and *M. baffinensis* are indicated by arrows.

Supplementary Figure 3. Maximum-likelihood phylogenetic tree inferred from nuclear 18S rRNA sequences of Mamiellophyceae. *Monomastix opisthostigma* was used as an outgroup. Solid dots correspond to nodes with significant support (> 0.8) for ML analysis and Bayesian analysis (>0.95). Empty dots correspond to nodes with non-significant support for either ML or Bayesian analysis, or both. GenBank accessions of the 18S rRNA sequences shown after the species name.

Supplementary Figure 4. Intramolecular folding pattern of the ITS2 molecule of *Mantoniella* (RCC2288, RCC2285, RCC2497 and RCC5418). The four major helices are labeled as Helix I – Helix IV. Blue dots represent either CBCs or hCBCs. Non-CBCs (N – N ↔ N × N) are represented in orange.

Supplementary Figure 5. Molecular signatures of *Mantoniella* species revealed by comparison of ITS2 secondary structures within Mamiellaceae. Signatures in of Helix III are shown in (**A**) and Helix IV in (**B**). The conserved base pairs among the different groups are numbered. CBCs and hCBCs are highlighted by solid and dotted arrows, respectively. Hypervariable positions are marked by an asterisk (*). Ellipsis (…) represent the other clades and species of *Micromonas*. The YRRY (pyrimidine-purine-pyrimidine) motif on the 5’ side arm of Helix III is shown in bold black. Single nucleotide substitutions are shown by grey nucleotides. Identified homoplasious changes are shown as parallelisms and reversals.

## Supplementary Tables

Supplementary Table 1. Primers and PCR conditions used in this study. Abbreviations: fwd – forward, rev. – reverse, Temp. – Temperature.

Supplementary Table 2. Metabarcoding datasets of the 18S rRNA gene analysed in this study for the presence of *M. beaufortii* and *M. baffinensis* signatures.

Supplementary Table 3. Morphological characters in Mamiellophyceae species.

Supplementary Table 4. Pigment composition of *M. beaufortii* (RCC2288) compared to a selection of green algae. Values are shown as a ratio of pigment to Chl *a* concentration and percent contribution to total carotenoids (in italics). See Supplementary Table 5 for the full names of the pigments.

Supplementary Table 5. Pigments analyzed in this study. LOD, limit of detection.

## Supplementary Material

Supplementary Material 1: Video microscopy of RCC2288 swimming behavior (https://youtu.be/CGKNxzfGUvQ).

Supplementary Material 2: Video microscopy of RCC2497 swimming behavior (https://youtu.be/rRNuk5Lx7Aw).

Supplementary Material 3: Video microscopy of RCC5418 swimming behavior (https://youtu.be/xoxCEl1cv4Q).

## Abbreviations

rRNA: ribosomal RNA
ITS2: internal transcribed spacer 2
CBC: compensatory base change
hCBC: hemi-CBC
TEM: transmission electron microscopy

