## Supplementary figures and images for "*Mantoniella Beaufortii* and *Mantoniella Baffinensis* sp. nov. (Mamiellales, Mamiellophyceae), Two New Green Algal Species from the High Arctic"

### Supplementary Figure 1

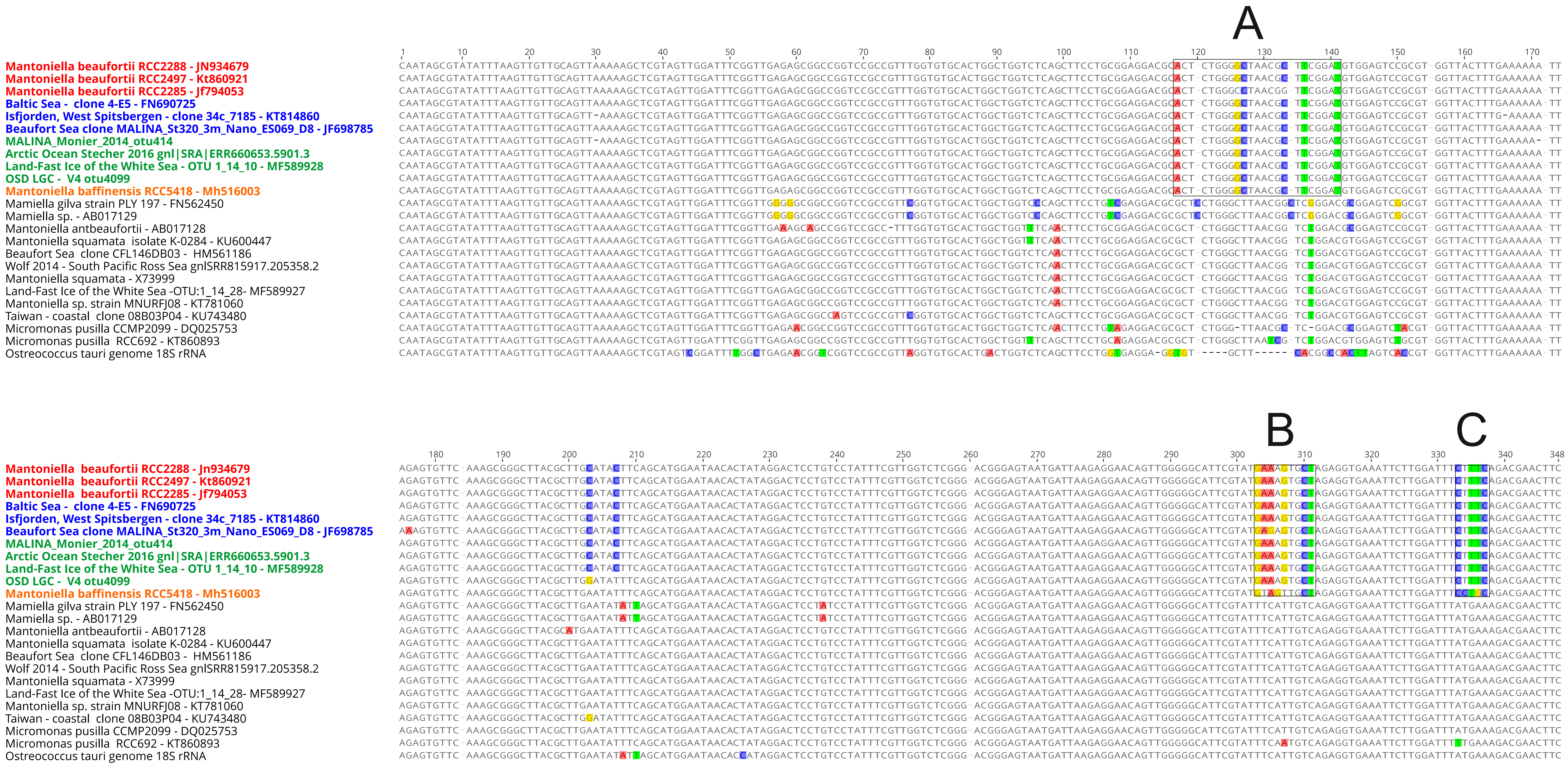

### Supplementary Figure 2

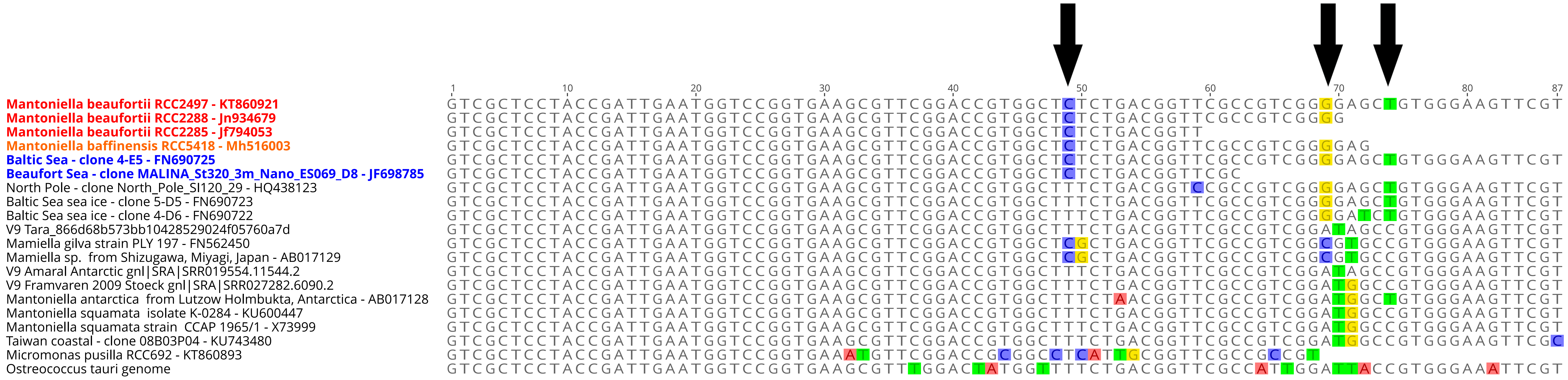

### Supplementary Figure 3

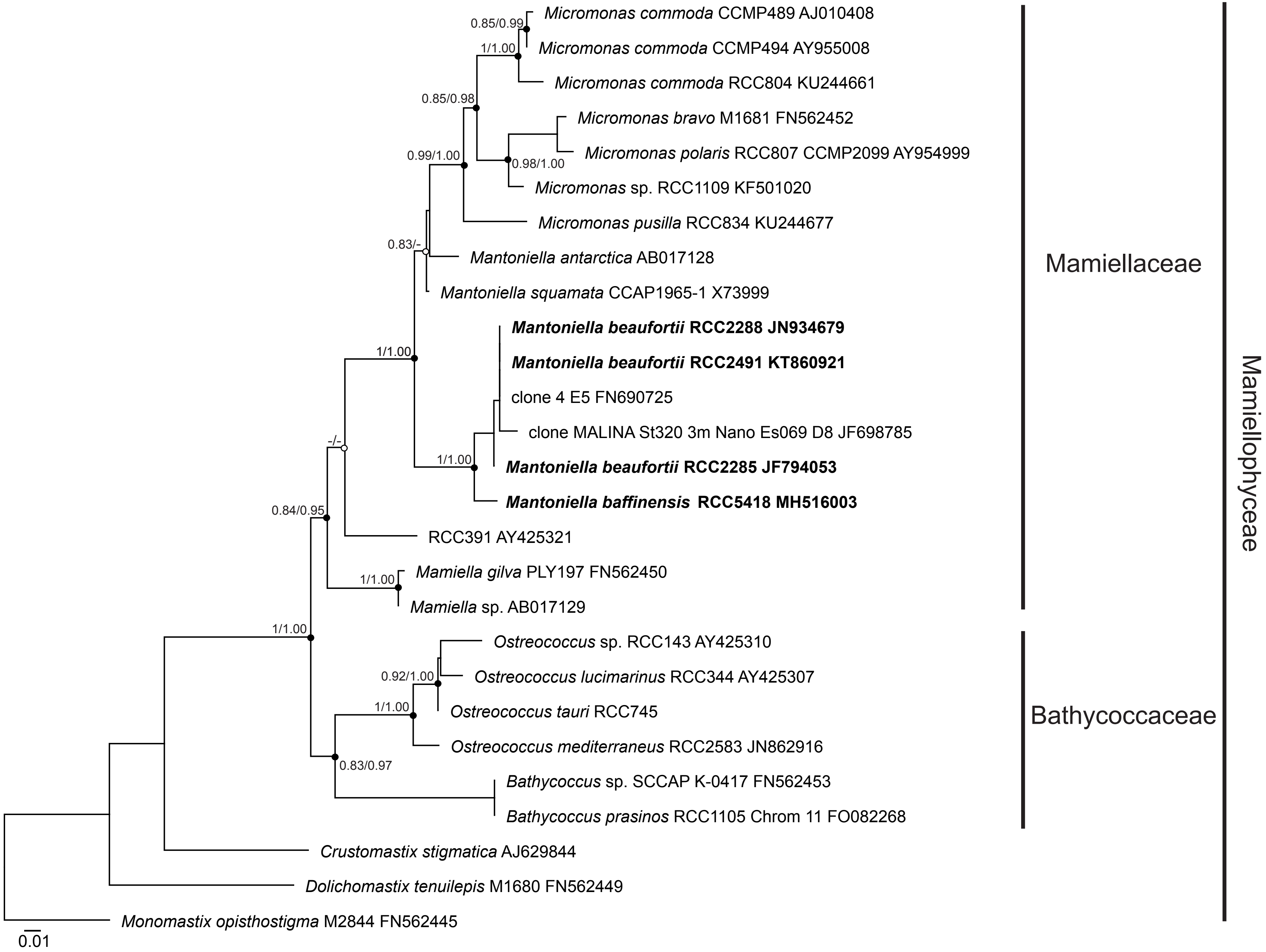

### Supplementary Figure 4

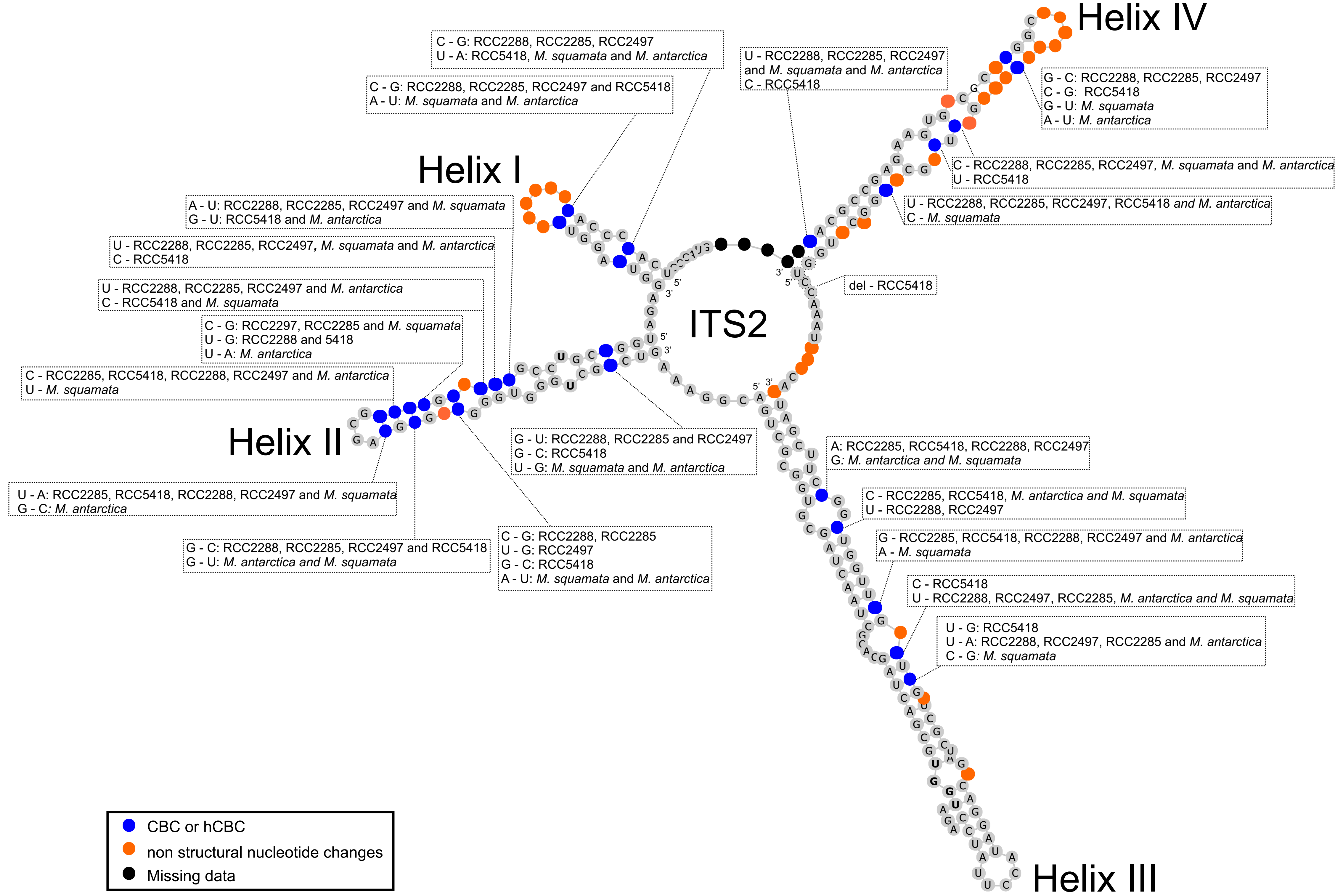

### Supplementary Figure 5A

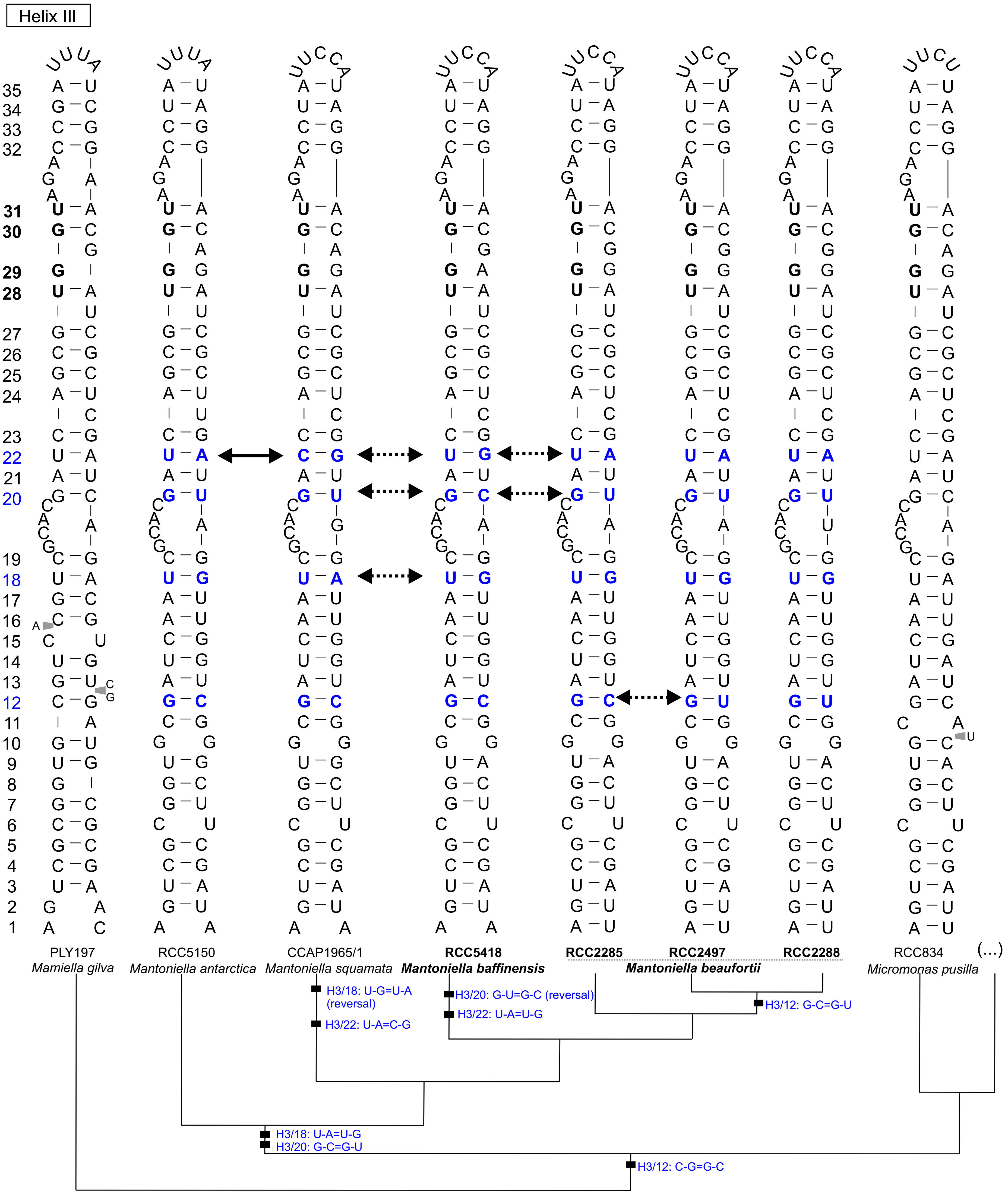

### Supplementary Figure 5B

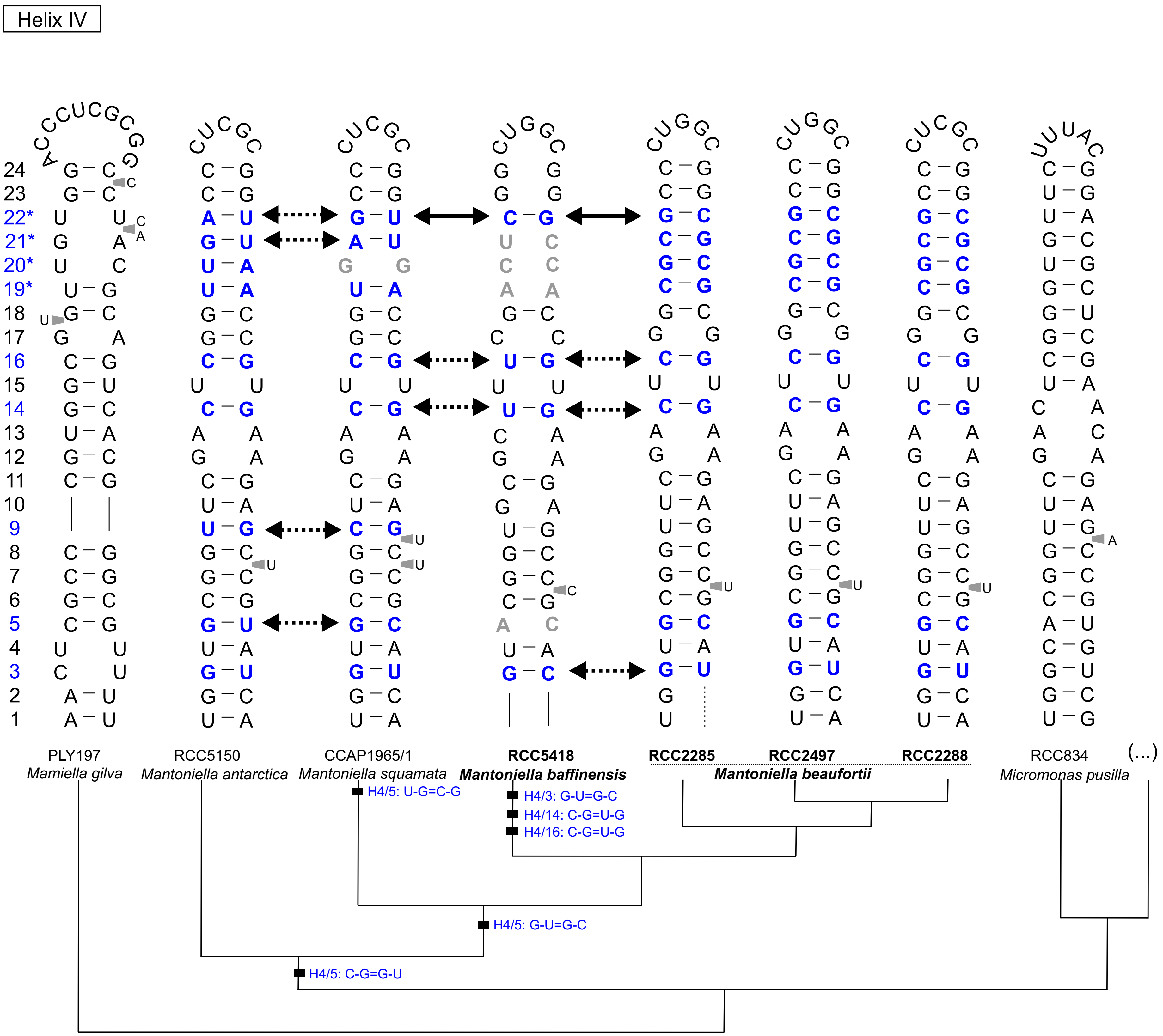
